# Effects of heavy water on protein dynamics studied by molecular dynamics simulation: Focusing on dynamical parameters obtained by quasi-elastic neutron scattering

**DOI:** 10.1101/2022.09.08.507213

**Authors:** Tatsuhito Matsuo

**Affiliations:** Institute for Quantum Life Science, National Institutes for Quantum Science and Technology, 2-4 Shirakata, Tokai, Ibaraki, 319-1106, Japan

**Author notes:** Corresponding author: Tatsuhito Matsuo, Ph.D.

## Abstract

Quasi-elastic neutron scattering (QENS) is a powerful technique to study protein dynamics. In general, QENS measurements are carried out in D_2_O solvent whereas functional studies of proteins are conducted in H_2_O solvent. Therefore, to link the QENS studies with the functional studies and then to understand the molecular basis of protein functions in detail, it is important to investigate the effects of solvent isotopic change on dynamical parameters obtained by QENS. For this purpose, in this study, MD simulations were carried out on hen egg white lysozyme, a well-folded and characterized protein, in H_2_O and in D_2_O. The dynamical parameters were extracted from the QENS spectra calculated from the MD trajectories. It was found that isotopic effects depend on energy resolutions and that at the energy resolutions that recent QENS studies often employ, the local dynamical behavior of proteins characterized in D_2_O more or less reflects that in H_2_O.

## Introduction

Proteins are bio-macromolecules essential for life, involved in diverse cellular functions by catalyzing biochemical reactions or playing a structural role to maintain the internal architecture of cells or organelles [1]. They are dynamic objects and are constantly fluctuating under the influence of thermal energy of water molecules inside the cell. Since utilization of the thermal fluctuation is known to be essential for protein functions [2], it is of critical importance to establish a relationship between molecular dynamics and protein functions for understanding the molecular mechanism of protein function.

In general, biochemical and biophysical studies focusing on functional aspects of proteins such as kinetics, activity, and interactions, are carried out in H_2_O-based buffer. On the other hand, quasielastic neutron scattering (QENS), which is a powerful tool to investigate intramolecular motions, is typically used for proteins hydrated by D_2_O or dissolved into D_2_O-based buffer [3], i.e. in different solvent conditions from those of functional studies. QENS has widely been used to investigate the physical responses of protein molecules to perturbations such as temperature change and change in hydration level within the framework of solid state physics and molecular physics since the pioneering work in 1989 by Doster et al. [3–6]. Several studies have focused on the dynamics-function relationship of proteins such as bacteriorhodopsin [7–10] and those in photosystem II membrane fragments [11,12]. In addition to this, a new research trend focusing on protein dynamics in light of functional aberration of proteins, i.e., human diseases and disorders, has emerged during the last decade ([13] and references therein). This trend, which is associated with molecular physiology and pathology, emphasizes more than ever the importance to interpret QENS data in association with protein functions, which are determined in light water environments. It is, therefore, essential to understand how different the dynamical parameters describing the molecular dynamics of proteins observed by QENS are between in H_2_O and in D_2_O in order to link QENS studies and functional studies and thus provide ultimately atomic-level physical interpretation of protein functions.

So far, some experimental and theoretical studies have focused on the effects of the deuteration of solvent water molecules on protein dynamics [14–17]. It has been shown by molecular dynamics (MD) simulation on myoglobin molecules partially hydrated with H_2_O or D_2_O that the mean square fluctuation was smaller for the latter than for the former at 300 K [14]. Another MD simulation study has found that copper plastocyanin in D_2_O increased the number of intra-protein hydrogen bonds compared with that in H_2_O [15]. Experimentally, it has been reported that there is a tendency that rigidification of protein molecules occurs in D_2_O compared with in H_2_O [16] and that hydrodynamic radii of RNase A and ubiquitin decrease by 2–5% by the solvent isotopic change [17]. These studies suggest a repressive effect of D_2_O on protein flexibility and have advanced our understanding of physicochemical properties of proteins in H_2_O and in D_2_O. However, little is known about the solvent isotopic effects on dynamical parameters that can be extracted from QENS measurements.

QENS permits to characterize molecular dynamics of a target protein at the sub-nanosecond timescale and the Å length-scale. The measured QENS spectra are dominated by scattering of hydrogen (H) atoms in the samples because the incoherent neutron scattering cross-section of H atoms is more than 40 times larger than any other atoms found in proteins and deuterium (D) atoms. Since the number of H atoms account for about half of the total number of atoms constituting protein molecules and they are distributed quasi-uniformly in space, dynamical information extracted from the QENS spectra reflect the motions of H atoms averaged over all the H atoms in proteins. To be precise, since QENS studies on proteins generally employ D_2_O solution samples to minimize the scattering contribution of solvent to the measured QENS spectra, labile H atoms of proteins, which are bound to N and O, are exchanged with D atoms of the solvent. Thus, the motions of non-exchangeable H atoms are characterized by QENS in the case of proteins in D_2_O solution.

Here, in order to study possible effects of isotopic change from H to D in the solvent on dynamical parameters obtained by QENS, MD simulation was employed for hen egg white lysozyme (HEWL), a well-folded and characterized protein, in H_2_O and in D_2_O using the latest force field for heavy water [18]. It was found that isotopic effects depend on energy resolutions and that at the energy resolutions that recent QENS studies often employ, the local dynamical behavior of proteins characterized in D_2_O more or less reflects that in H_2_O.

## Materials and methods

### Molecular dynamics simulation

Molecular dynamics (MD) simulation was performed using the GROMACS software (version 2018.3) [19] with the AMBER03 force field [20]. As light and heavy water models, the TIP3P model [21] and the latest heavy water model, TIP3P-HW [18], were employed, respectively. A hen egg white lysozyme (HEWL) molecule (PDB ID: 2AKI; Fig. 1 (a)) was solvated in a cubic box such that the protein was placed at ≥ 10 Å from the edges of the simulation box. To neutralize the charges of the entire system, counter ions were added. van der Waals force and electrostatic interactions were cut off at 8.0 Å [20]. The temperature was set to be 300 K and the pressure was 1 atm.

**Figure 1.**
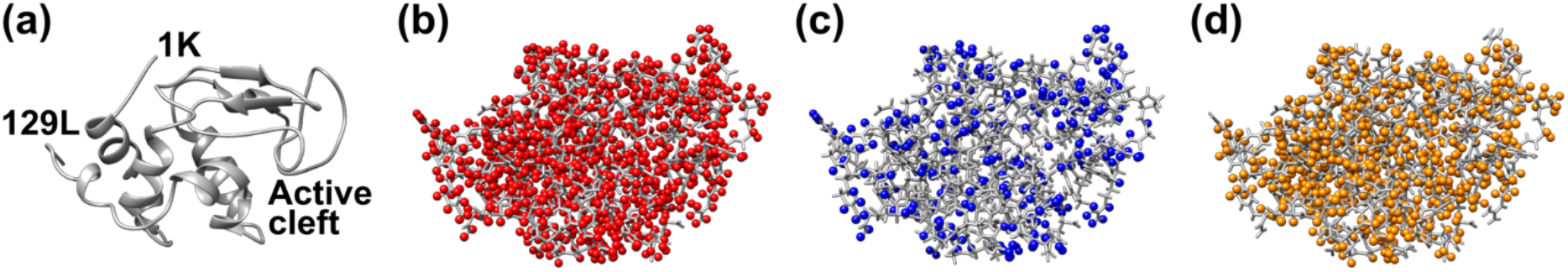
Atomic structure of a HEWL monomer. (a): The HEWL monomer is displayed in ribbon representation. The N- and C-terminal residues are also shown. (b): All the hydrogen atoms in HEWL are shown in red spheres. (c, d): Exchangeable (labile) hydrogen atoms (c) and non-exchangeable hydrogen atoms (d) are shown in blue and orange spheres, respectively.

In the simulation, three samples were employed: HEWL in H_2_O (LysH), HEWL in D_2_O with no exchange in labile hydrogen atoms, which is a quasi-”in D_2_O” state (LysQD), and HEWL in D_2_O with H/D exchange of labile hydrogen atoms (LysD). For reference, the spatial distributions of all, non-exchangeable and exchangeable hydrogen atoms are shown in Fig. 1 (b–d). The H/D exchange was done by simply doubling the mass of the corresponding hydrogen atoms as in previous studies [14,15,22]. Note that whereas the simple mass substitution fails to reproduce an experimentally-observed molecular behavior occurring much longer time scale, e.g., 400 ns [23], there are not significant errors up to 50 ns from the start of the simulation. The current simulation time of 15 ns (see below) is thus short enough not to induce large errors in the dynamical behavior of protein atoms and long enough to estimate dynamical parameters obtained by QENS. Comparison of the results from LysH and LysQD allows to observe the pure effects of solvent deuteration while comparison of the results from LysH and LysD allows to observe the “real” isotopic effects on protein dynamics taking H/D exchange of labile hydrogen atoms into account.

For each condition, the system was energy-minimized by the steepest descent method. NVT and NPT equilibration was subsequently carried out for 100 ps each with a time step of 2 fs. The resultant structure was subjected to a production run for 15 ns, which was conducted in triplicate. The trajectories were saved every 2 ps. For each trajectory, protein molecules were aligned to remove translational and rotational motions of the whole molecule. The resultant trajectories were used for further analysis. The density of solvent was 1004.7, 1104.7, and 1105.9 kg/m^3^ on average for LysH, LysQD, and LysD, respectively. Latter two values are in agreement with 1104.0 at 300 K estimated based on the equation of state for D_2_O [24]. Moreover, the translational diffusion coefficient (D_T_) of the entire protein molecule, which was estimated from the mean square displacement using the GROMACS command “gmx msd” for the MD trajectories before translational and rotational alignments, was 1.96 × 10^−6^, 1.65 × 10^−6^, and 1.52 × 10^−6^ cm^2^/s for LysH, LysQD, and LysD, respectively.

Smaller D_T_ values for LysQD and LysD than LysH are consistent with the larger solvent viscosity of D_2_O than that of H_2_O [17]. These results confirm the use of D_2_O as solvent in the simulations of LysQD and LysD.

### Calculation of intermediate scattering functions and dynamic structure factors from MD trajectories

The spectra measured by QENS are described by the incoherent dynamic structure factor 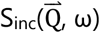, where 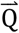, and ω are the momentum transfer and the energy transfer of neutrons, respectively. In order to calculate 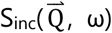 from a MD trajectory, it is straightforward to calculate first the incoherent intermediate scattering function 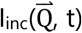, which is written as

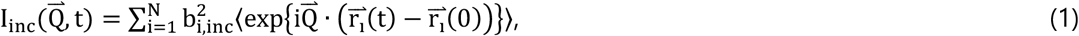

where N is the total number of atoms contained in a protein and b_i,inc_ is the incoherent scattering length of the i-th atom. 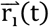 is the position vector of the i-th atom at time t, which is given by a MD trajectory, and 〈 〉 denotes the thermal average. In the case of solution samples, 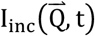 is orientationally averaged since protein molecules take various orientations, and the orientation average and the thermal average are independent. This leads to the following equation:

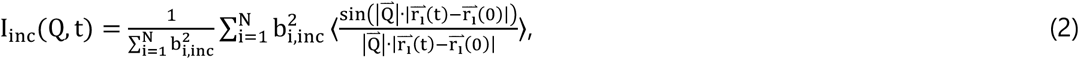

where the pre-factor is a normalization factor (I_inc_(Q, 0) = 1). The b_i,inc_ value of 25.217 was used for hydrogen atoms, while other atoms were omitted from the calculation since the b_i,inc_ values of these atoms are negligible [25]. It was confirmed that additional inclusion of the contribution of deuterium atoms (b_i,inc_ = 4.033) that replace the exchangeable H atoms in LysD gave negligible effects on the I_inc_(Q, t) profiles (data not shown).

The dynamic structure factor S_inc_(Q, ω) is described as the time Fourier transform of I_inc_(Q,t),

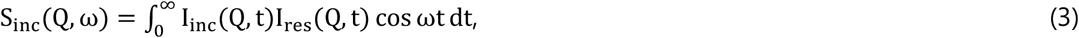

where I_res_(Q, t) is an instrumental resolution function, which defines the time-window of observation. In this study, this function was represented by a Gaussian function as,

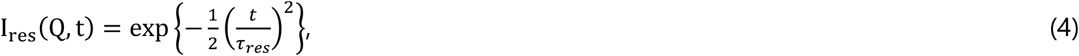

where *τ_res_* is the resolution time. As *τ_res_* values, 10, 50, and 100 [ps] were employed in the analysis of S_inc_(Q, ω), which correspond to the energy resolutions (FWHM) of 77, 15, and 7.7 [μeV], respectively. These values are within the range employed in many QENS studies.

### Analysis of the intermediate scattering function

In order to extract dynamical information using I_inc_(Q,t) calculated from a MD trajectory, I_inc_(Q,t) was fitted by the following equation [25]:

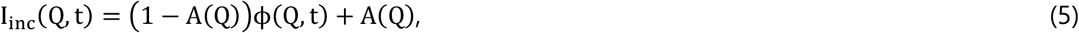

where the first and the second terms represent the quasi-elastic and the elastic component, respectively. A(Q) is the elastic incoherent structure factor (EISF). The time evolution of I_inc_(Q,t) is characterized by ϕ(Q, t). As was demonstrated before [26], it was not possible to fit the calculated I_inc_(Q,t) with a single exponential function. ϕ(Q, t) was thus represented by the so-called Kohlrausch-Williams-Watts stretched exponential function [27],

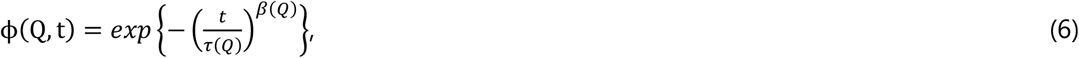

where *τ*(*Q*) is the relaxation time and *β*(*Q*) is a measure of the distribution of the relaxation times. As easily found, *β*(*Q*) = 1 retrieves a single exponential function.

### Analysis of the dynamic structure factor

Dynamical parameters obtained by QENS measurements were estimated by fitting the calculated S_inc_(Q, ω) with the following phenomenological equation:

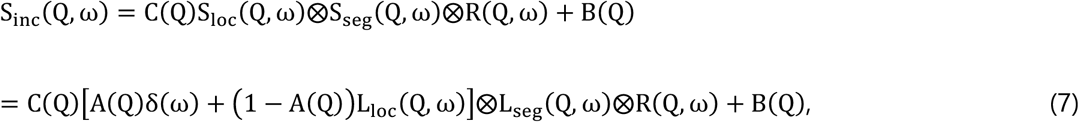

where S_loc_(Q, ω) represents the contribution from local atomic motions in the protein. S_seg_(Q, ω) denotes the contribution from motions occurring at slower time scale such as backbone and segmental motions. Since the protein molecules have been aligned to remove translational and rotational movements of the entire molecule as mentioned above, these motions are negligible in the calculated QENS spectra and hence their contributions are not included in Eq. 7. C(Q) is a scaling factor between the fit and the calculated spectra. R(Q, ω) and B(Q) are a resolution function and a background, respectively. L_loc_(Q,ω) is a Lorentzian function describing local atomic motions and in the form of

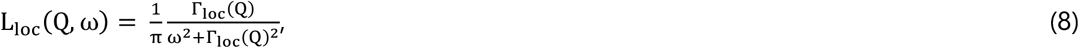

where Γ_loc_(Q) is the half width at the half maximum (HWHM) of the Lorentzian. L_seg_(Q,ω) also takes the same form with the HWHM of Γ_seg_(Q). The term including δ(ω) is the elastic component and the term including L_loc_(Q,ω) is the quasi-elastic component.

Q^2^-dependence of Γ_loc_ (Q) and Γ_seg_(Q) provides the information on the nature of the corresponding diffusive motions. It was found that Γ_loc_ (Q) and Γ_seg_(Q) obtained by the fitting of the calculated QENS spectra using Eq. 7 are approximated by the jump-diffusion model [28]:

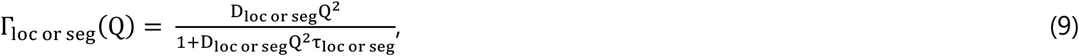

where τ_loc_ and τ_seg_ are the residence times of local motions and segmental motions, respectively. Similarly, D_loc_ and D_seg_ are the jump-diffusion coefficients of local motions and segmental motions, respectively.

Q-dependence of the EISF A(Q) provides the information on the geometry of motions. In this study, the EISF was fitted using either of the following equations, which describe a diffusion-inside-a-sphere model [29], as appropriate:

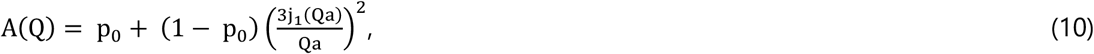

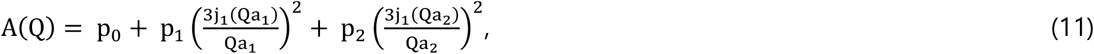

where p_0_ is the immobile fraction whose motions are too slow to be observed within the energy resolution employed. In Eq. 10, (1 – p_0_) denotes a fraction of atoms observed at the energy resolution employed and such atoms undergo diffusive motions within spheres of a radius of a. j_1_(x) is the first-order spherical Bessel function of the first kind. In Eq. 11, p_1_ and p_2_ denote fractions of atoms moving within spheres of the radii of a_1_ and a_2_, respectively. p_0_, p_1_, and p_2_ satisfy the condition p_0_ + p_1_ + p_2_ = 1.

### Statistical test

Comparisons of dynamical parameters were carried out by the two-paired Student’s t-test. In the case of comparisons of three parameters, paired comparisons need to be made three times, which increases the significant level (p) from 0.05 to 0.14. Therefore, in this case, the so-called Bonferroni correction was employed [30], where the value of the significant level was divided by three to compensate for this effect.

## Results and discussion

### MD trajectories

Fig. 2 shows the time course of the RMSD values calculated from the MD trajectories. It was found that the RMSD values are stable around 0.12 Å after 5 ns of the start of the simulations. Therefore, MD trajectories in the time range of 5–15 ns were used to calculate neutron scattering functions as described in the following sections. Radii of gyration of HEWL averaged over three trajectories were 14.2 ± 0.1 [Å], 14.2 ± 0.1 [Å], and 14.3 ± 0.1 [Å] for LysH, LysQD, and LysD, respectively. These values are consistent with 14.12 ± 0.10 [Å] reported in a previous study [31] and suggest that the overall structure of HEWL is not affected by solvent exchange nor H/D exchange of labile H atoms within the time regime employed.

**Figure 2.**
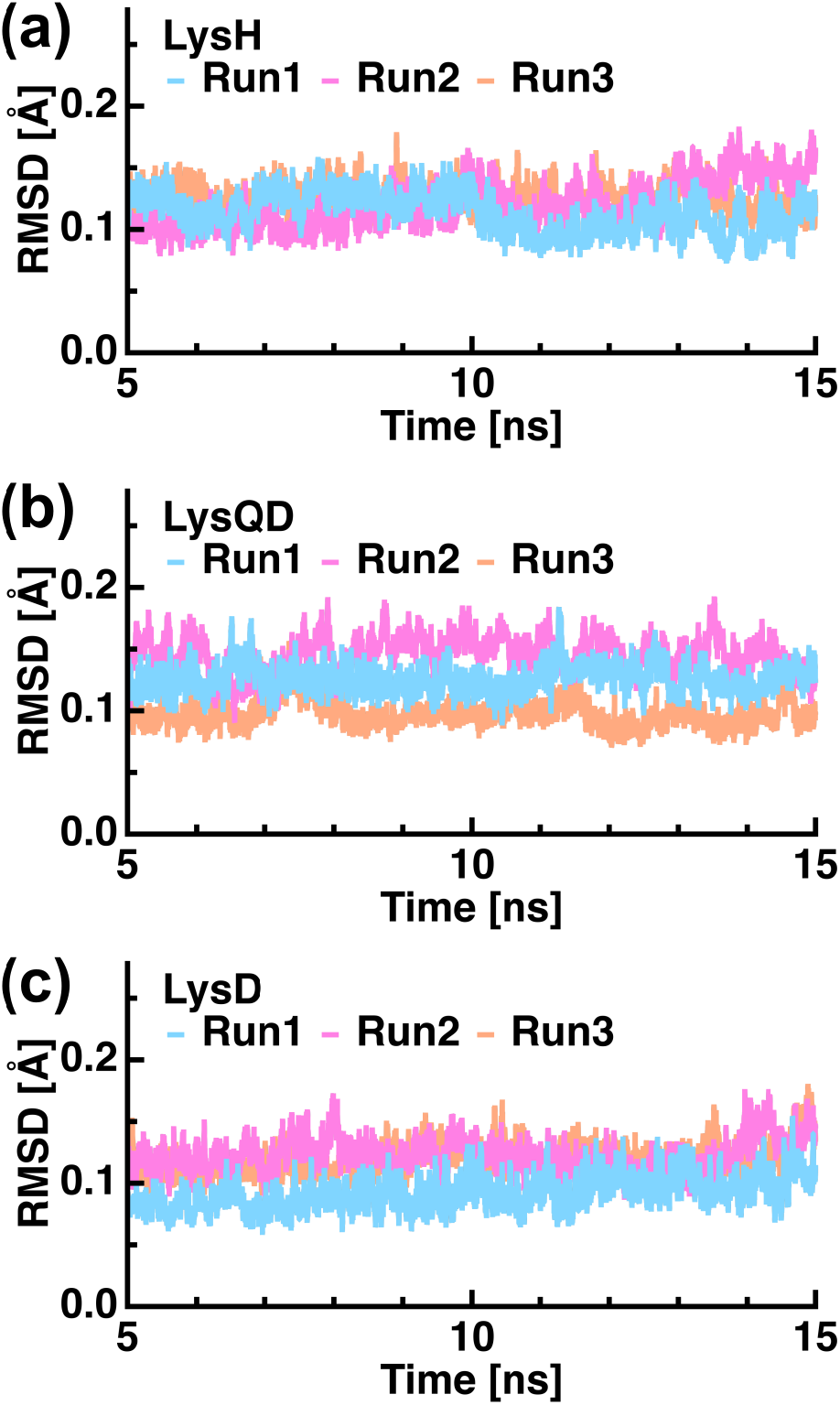
RMSD values as a function of time. Those between 5 ns and 15 ns, which were used for subsequent analyses are shown. In each graph, the RMSD values of the 1^st^, 2^nd^, and the 3^rd^ runs are shown together.

### Incoherent Intermediate scattering function I_inc_(Q, t)

In typical QENS experiments, D_2_O solution samples of proteins are employed as specimens. Because of H/D exchange between labile H atoms and solvent D atoms, QENS spectra arise from the contribution of non-exchangeable H atoms (NEX) in the proteins. In order to directly compare the isotopic effects on QENS-related dynamical parameters, incoherent intermediate scattering functions I_inc_(Q, t) and incoherent dynamic structure factors S_inc_(Q, ω) were evaluated from NEX atoms of LysH, LysQD, and LysD.

The I_inc_(Q, t) curves of NEX atoms of LysH, LysQD, and LysD calculated using Eq. 2 are shown in Fig. 3 (a), (b), and (c), respectively. These curves were the averages of three I_inc_(Q, t) curves calculated from each MD trajectory and were fitted quite well by Eqs. 5 and 6, providing dynamical parameters shown in Fig. 3 (d-f). EISF(Q) was fitted by Eq. 11, which contains two populations of atoms undergoing diffusive motions within spheres with different radii. The resultant geometrical parameters were tabulated in Table 1. It was found that the immobile fraction of atoms (p_1_) is larger for NEX-LysQD (p < 0.05) and NEX-LysD (p = 0.10), and the fraction of atoms moving with larger radii (p_1_) is smaller for NEX-LysD (p < 0.05) than NEX-LysH while there were no significant differences in other parameters. These results suggest that the mobility of more atoms is slowed down in D_2_O than in H_2_O. The increase in p_0_ and decrease in p_1_ without changing the amplitudes observed for NEX-LysD result in the averaged amplitude smaller than that of NEX-LysH. This is consistent with a previous MD study which found that the mean-square fluctuation of myoglobin in a hydrated powder state decreases in D_2_O compared with in H_2_O [14].

**Figure 3.**
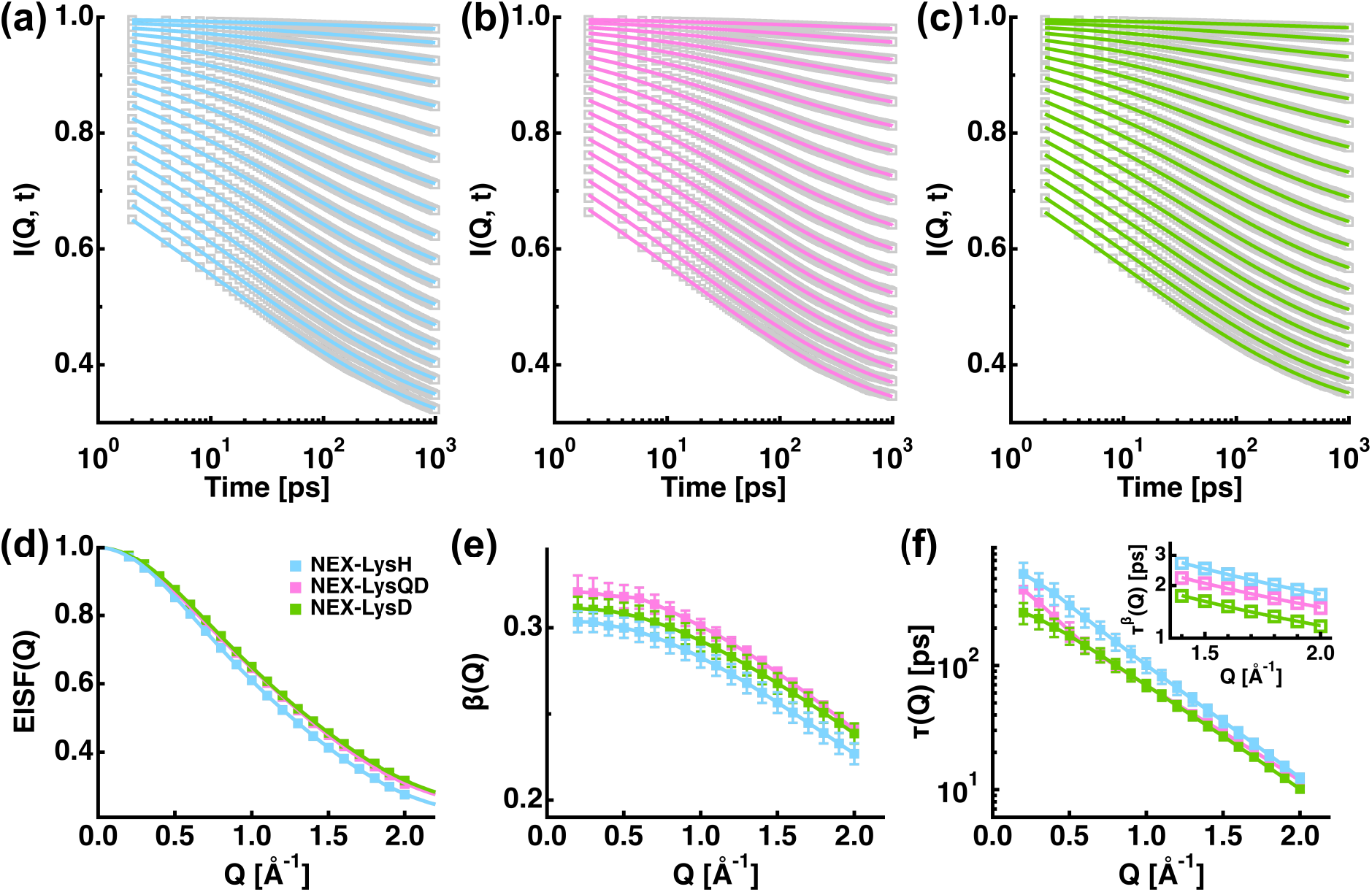
Incoherent intermediate scattering functions I_inc_(Q, t) calculated from MD trajectories for non-exchangeable hydrogen atoms (NEX) of LysH (a), LysQD (b), and LysD (c). I_inc_(Q, t) is normalized to 1.0 at t = 0. Grey squares represent the calculated I_inc_(Q, t) values and solid lines represent the fitted ones using Eqs. 5 and 6 in the main text. From top to bottom, the corresponding Q values are from 0.2 [Å^−1^] to 2.0 [Å^−1^] with the interval of 0.1 [Å^−1^]. (d), (e), and (f) show the Q-dependence of EISF(Q), β(Q), and τ(Q), respectively, which were obtained from the fitting in (a–c). The inset of (f) shows the fitting results by Eq. 12. The curves are shifted vertically for clarity. Error bars are within symbols if not shown.

**Table 1.** Summary of parameters describing the geometry of motions obtained from EISF(Q). The values in parentheses are errors associated with the fitting.

| | $p_0$ | $p_1$ | $a_1$ [Å] | $p_2$ | $a_2$ [Å] |
| --- | --- | --- | --- | --- | --- |
| NEX-LysH | 0.213 (0.006) | 0.141 (0.007) | 3.70 (0.08) | 0.646 (0.009) | 1.55 (0.02) |
| NEX-LysQD | 0.236 (0.006) | 0.125 (0.006) | 3.42 (0.06) | 0.639 (0.008) | 1.48 (0.02) |
| NEX-LysD | 0.236 (0.008) | 0.107 (0.008) | 3.9 (0.1) | 0.66 (0.01) | 1.52 (0.03) |

As shown in Fig. 3 (e), the stretch factor β(Q) decreased from 0.30 to 0.23 for NEX-LysH, from 0.32 to 0.24 for NEX-LysQD, and from 0.31 to 0.24 for NEX-LysD as Q increased. As easily found from Eq. 6, non-exponentiality of relaxation behavior becomes greater as the β(Q) value deviates from 1.0. The current results imply that there is a tendency to reduce the non-exponentiality of local relaxational motions in D_2_O compared with in H_2_O. As shown in Fig. 3 (f), the relaxation times τ(Q) of all the three samples showed a decrease as Q increased, followed by a power-law behavior in the Q-region of 1.4–2.0 Å^−1^. The τ^β^(Q) values in this region, as shown in the inset of Fig. 3 (f), were fitted by

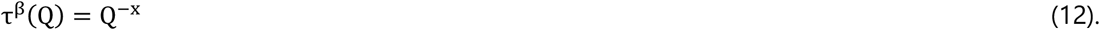

The resultant x values were 1.17 ± 0.02, 1.10 ± 0.02, and 1.11 ± 0.02 for NEX-LysH, NEX-LysQD, and NEX-LysD, respectively. These values provide a measure of homogeneity (or inhomogeneity) of relaxational behavior of atomic motions: The condition τ(Q) ~ Q^−2^ is satisfied when the stretched exponential behavior of I_inc_(Q, t) arises from an ensemble of exponential functions with various relaxation times (“Homogeneous” scenario) [32]. On the other hand, when the stretched exponential behavior arises from an ensemble of stretched exponential functions with the identical relaxation time (“Heterogeneous”scenario), the condition τ(Q) ~ Q^−2/β^ is satisfied. The values of 2/β in the present data are between 7.62 and 8.82 (NEX-LysH), between 7.15 and 8.30 (NEX-LysQD), and between 7.32 and 8.38 (NEX-LysD) while the values of x/β are between 4.45 and 5.14 (NEX-LysH), between 4.07 and 4.66 (NEX-LysQD), and between 3.93 and 4.57 (NEX-LysD). This comparison indicates that the relaxational behavior of protein atoms is intermediate between the homogeneous and heterogeneous scenarios, and it is a little bit closer to the homogeneous scenario, which is in line with a previous study [26].

I_inc_(Q, t) of exchangeable H atoms showed oscillations at longer timescales (data not shown) and could not be fitted with Eq. 5. This phenomenon is previously observed [26] and is ascribed to long-time statistical errors due to much smaller number of exchangeable H atoms than NEX atoms. Therefore, analysis of exchangeable H atoms was not carried out here.

### Incoherent dynamic structure factor S_inc_(Q, ω)

Next, to compare dynamical parameters obtained from QENS measurements, incoherent dynamic structure factors S_inc_(Q, ω) of NEX atoms were calculated using Eq. 3 with resolution times (*τ_res_*) of 10, 50, and 100 [ps]. Since S_inc_(Q, ω) is the physical quantity measured directly in real QENS measurements, the simulated QENS spectra calculated from each of three MD trajectories were analyzed without averaging to study the dynamical parameters in detail. The resultant QENS spectra were fitted with Eq. 7 and fitting examples are shown in Fig. 4. Note that in the case of *τ_res_* = 10 [ps], inclusion of the L_seg_(Q, ω) term in Eq. 7 did not converge the fitting, implying that segmental motions, which occur more slowly than local motions, are not resolved at this time resolution. Therefore, broadening of the elastic component is seen only in the cases of *τ_res_* = 50 and 100 [ps] (Fig. 4 (b) and (c)).

**Figure 4.**
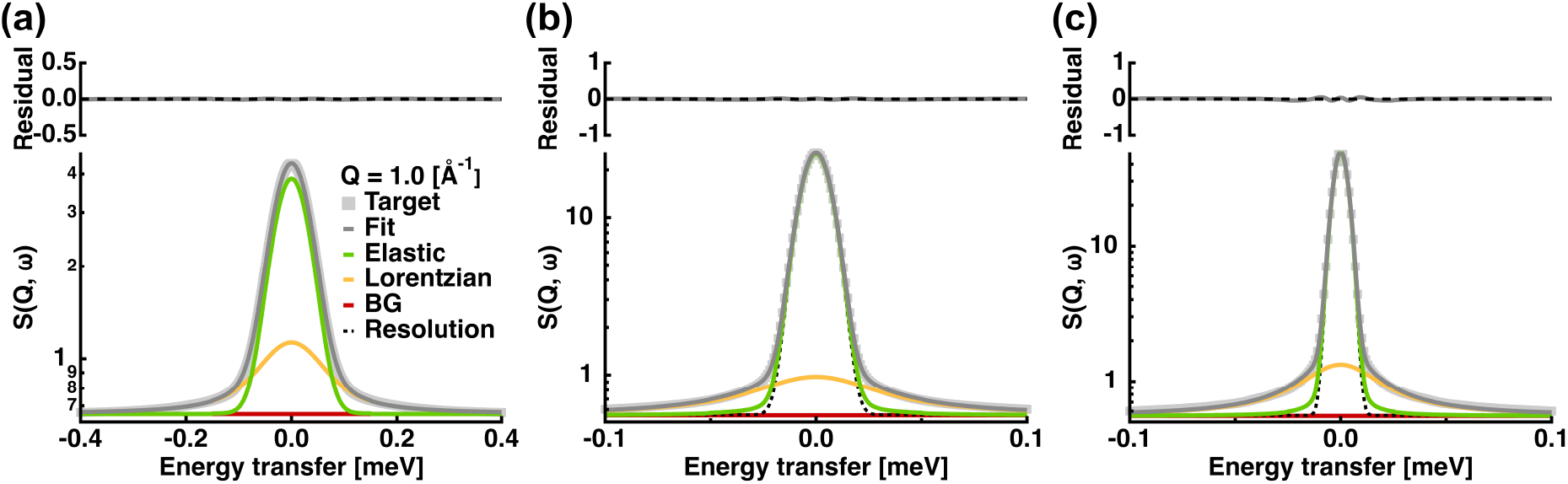
Examples of the fitting of the incoherent dynamic structure factor S_inc_(Q, ω) calculated from MD trajectories. Those with time resolutions of 10 ps (a), 50 ps (b), and 100 ps (c) of NEX-LysH at Q = 1.0 [Å^−1^] are shown. “Target” denotes S_inc_(Q, ω) calculated from MD trajectories. The upper panels are the residuals between the “Target” and the fitted curves. In (a), the resolution function is the same as the elastic component while broadening of the elastic component is seen in (b) and (c).

Fig. 5 summarizes the Q-(or Q^2^-) dependences of the EISFs and the widths of Lorentzian functions obtained from the fitting to the QENS spectra calculated from MD trajectories. The EISFs were able to be fitted by a diffusion-inside-a-sphere model with one population (Eq. 10) for all the time resolutions. Γ_loc_ (Q) at τ_res_ = 10 [ps] showed an asymptotic behavior as Q increased, suggesting a jump diffusion process. Γ_loc_(Q) at τ_res_ = 50 and 100 [ps] showed clearly more or less constant values at the low-Q region, indicating that the local motions can be described by a jump diffusion within a confined space. Γ_seg_(Q) observed only at τ_res_ = 50 and 100 [ps] also showed an asymptotic behavior with increase in Q, which is characteristic of a jump diffusion, suggesting that the discrete step of segmental motions is resolved at these slower time windows. Dynamical parameters describing the diffusive nature of motions were extracted by fitting of the Q^2^-dependences of Γ_loc_(Q) and Γ_seg_(Q) using Eq. 9. The extracted parameters, together with those obtained from the fitting of the EISFs, are summarized in Fig. 6. It was found that at *τ_res_* = 10 [ps], the D_loc_ value decreases and the τ_loc_ value increases by mere solvent change (p < 0.05), followed by non-significant change in these parameters by additional H/D exchange of labile H atoms. These results suggest that the mere effect of solvent change from light water to heavy water has large effects on local motions whereas D atoms that have replaced labile H atoms do not modulate the mobility of non-exchangeable atoms. Regarding the geometry of motions, there were no significant difference in the radius nor in the immobile fraction, implying that isotopic effects are manifested in the rates of local motions, not in their spatial features. This is in agreement with a physical property of D atoms where hydrogen bonding is strengthened by exchange from H to D [33], which restrains the kinetic aspect of atomic motions. Moreover, this is also in line with a previous finding that mobility of buried tryptophan residues in several proteins is slowed down in D_2_O compared with in H_2_O [16]. At *τ_res_* = 50 [ps], a significant increase in τ_loc_ by H/D exchange of labile H atoms is still observed while other parameters including segmental motions are not affected by isotopic exchange in solvent and/or labile H atoms. At *τ_res_* = 100 [ps], all the dynamical parameters were the same within errors. These results indicate that local motions observed in a short time window are sensitive to isotopic effects while motions observed in longer time windows are not. It has been shown that some proteins show enhanced propensity to aggregation in D_2_O [34–37]. This might arise from modulation of local interactions involving amino acid side chains in the interface between the protein molecules, not from larger-scale motions occurring at much slower timescales.

**Figure 5.**
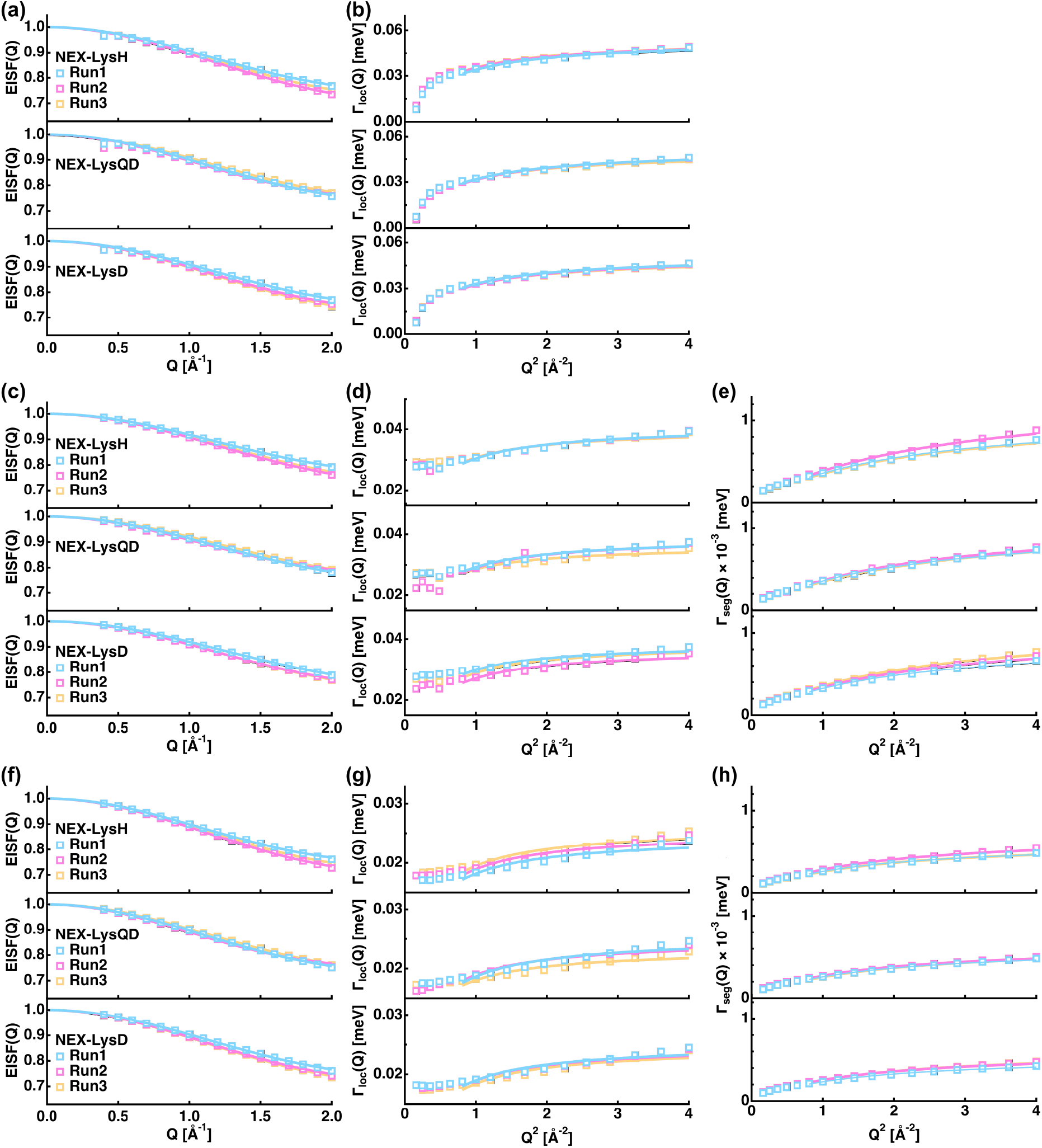
Summary of the Q-dependences of the EISFs (a, c, f) and the Q^2^-dependences of the Lorentzian widths describing local motions (b, d, g) and segmental motions (e, h) obtained from the fitting to the QENS spectra calculated from MD trajectories at the time resolutions of 10 ps (a–b), 50 ps (c–e), and 100 ps (f–h). Solid lines denote the fits to the spectra using Eqs. 8–10 in the main text.

**Figure 6.**
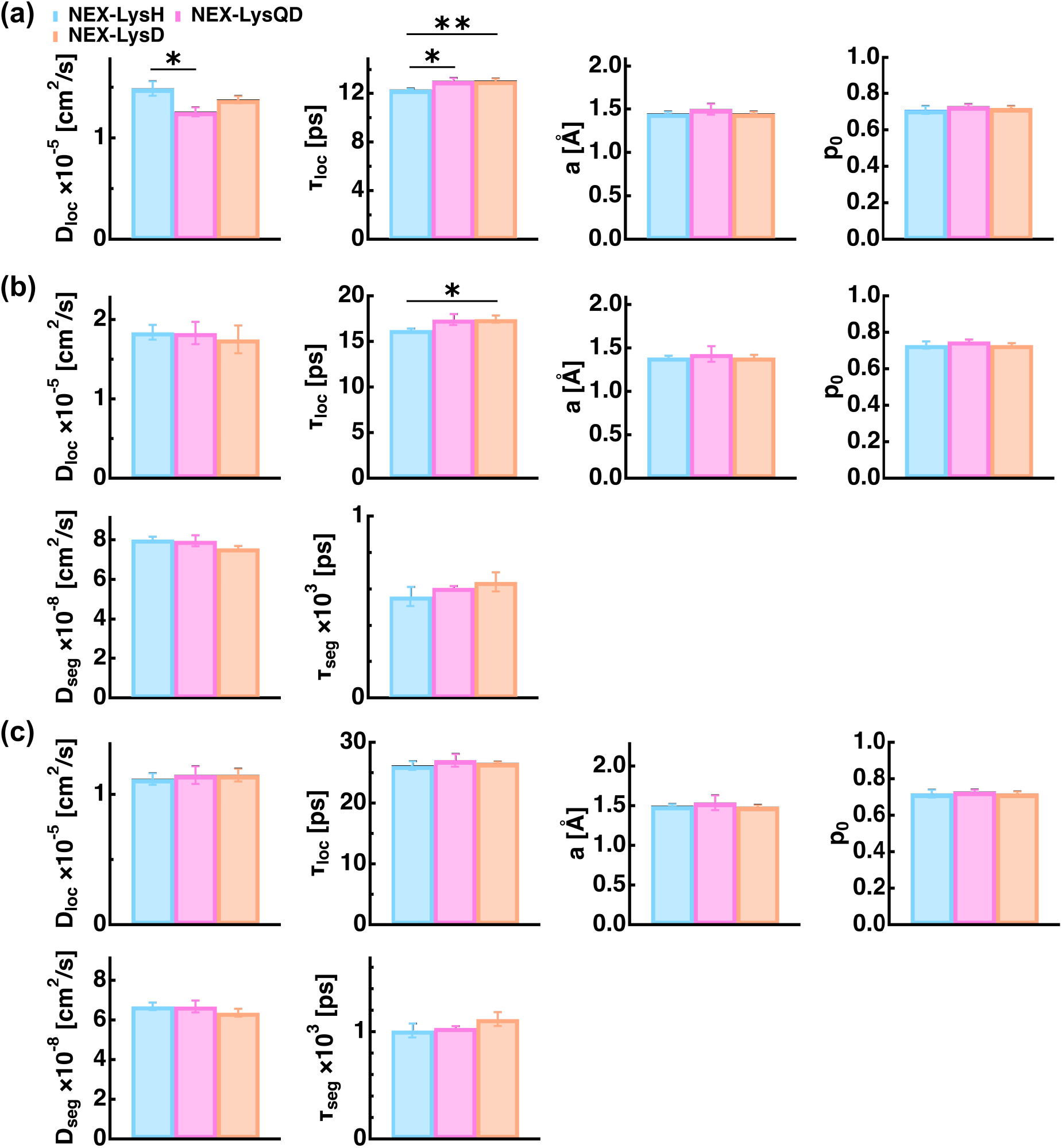
Comparison of dynamical parameters extracted from the EISFs and the Lorentzian widths shown in Fig. 5. For each parameter, the average and the standard deviation of the values obtained from three data sets (Run1–3) are shown. (a), (b), and (c) show the parameters obtained at the time resolutions of 10 ps, 50 ps, and 100 ps, respectively. Asterisks (*) indicate that the difference between the corresponding two parameters is statistically significant (p < 0.05 by Student’s t-test). The two asterisks in (a) denotes p < 0.001.

Recent QENS measurements on protein samples are often carried out at the energy resolutions of higher than 15 μeV (e.g. [38–40]). In these conditions, as seen in Fig. 6 (b) and (c), the majority of the dynamical parameters obtained by typical QENS measurements (NEX-LysD) is the same within errors as dynamical parameters that proteins show in H_2_O (NEX-LysH). Therefore, local dynamical behavior of proteins in D_2_O solvent resolved by QENS with high energy resolutions is considered to reflect that in light water in a good approximation. Since many biochemical and biophysical measurements to study kinetics and functions of proteins are carried out in light water environments, the present findings will serve as a foundation to link the dynamical features of proteins revealed by QENS with their biological functions, which will eventually lead to understanding of protein functions at the atomic level. Although a well-folded protein was employed in this study, it is conceivable that intrinsically disordered proteins respond to isotopic change differently. Future studies would be required to investigate this point in order to further deepen our knowledge of isotopic effects on protein dynamics.

### Conclusion

Intuitively, substitution of solvent or labile hydrogen atoms in proteins by heavier deuterium is expected to slow down the motions of protein molecules. However, the current study has revealed that the isotopic effects are time-resolution dependent and not all the dynamical parameters obtained by QENS are affected by such perturbations. At a narrow time window (*τ_res_* = 10 [ps]), the QENS parameters describing rates of motions such as the jump diffusion coefficient and the residence time were found to be sensitive to isotopic change and such parameters changed in the direction toward slower motions. However, these changes in dynamical parameters seem to become smaller or even disappear at longer time windows, which are the conditions often used in recent QENS studies. In these conditions, the dynamical behavior of target proteins in D_2_O solvent obtained by QENS is considered to reflect that in H_2_O, where functional and kinetic studies are generally carried out. Therefore, dynamical parameters extracted from QENS spectra will help understand the molecular basis of protein functions, even if the parameters are obtained in D_2_O environments.

## References

[1] B. Alberts, J. Alexander, L. Julian, R. Martin, R. Keith, W. Peter, Molecular biology of the cell, 5th ed., Garland Science, New York, 2008.

[2] G. Zaccai, How Soft Is a Protein? A Protein Dynamics Force Constant Measured by Neutron Scattering, Science (80-.)., 288 (2000) 1604–1607.

[3] F. Gabel, D. Bicout, U. Lehnert, M. Tehei, M. Weik, G. Zaccai, Protein dynamics studied by neutron scattering, Q. Rev. Biophys., 35 (2002) 327–367.

[4] W. Doster, S. Cusack, W. Petry, Dynamical transition of myoglobin revealed by inelastic neutron scattering, Nature, 337 (1989) 754–756.

[5] J.C. Smith, Protein dynamics: Comparison of simulations with inelastic neutron scattering experiments, Q. Rev. Biophys., 24 (1991) 227–291.

[6] V.G. Sakai, A. Arbe, Quasielastic neutron scattering in soft matter, Curr. Opin. Colloid Interface Sci., 14 (2009) 381–390.

[7] U. Lehnert, V. Reat, M. Weik, G. Zaccai, C. Pfister, Thermal motions in bacteriorhodopsin at different hydration levels studied by neutron scattering: correlation with kinetics and light-induced conformational changes, Biophys. J., 75 (1998) 1945–1952.

[8] J. Fitter, R.E. Lechner, N.A. Dencher, Picosecond molecular motions in bacteriorhodopsin from neutron scattering, Biophys. J., 73 (1997) 2126–2137.

[9] J. Fitter, R.E. Lechner, G. Buldt, N.A. Dencher, Internal molecular motions of bacteriorhodopsin: hydration-induced flexibility studied by quasielastic incoherent neutron scattering using oriented purple membranes, Proc Natl Acad Sci U S A, 93 (1996) 7600–7605.

[10] M. Ferrand, A.J. Dianoux, W. Petry, G. Zaccai, Thermal motions and function of bacteriorhodopsin in purple membranes: Effects of temperature and hydration studied by neutron scattering, Proc. Natl. Acad. Sci. U. S. A., 90 (1993) 9668–9672.

[11] J. Pieper, T. Hauss, A. Buchsteiner, G. Renger, The effect of hydration on protein flexibility in photosystem II of green plants studied by quasielastic neutron scattering, Eur. Biophys. J., 37 (2008) 657–663.

[12] G. Nagy, J. Pieper, S.B. Krumova, L. Kovács, M. Trapp, G. Garab, J. Peters, Dynamic properties of photosystem II membranes at physiological temperatures characterized by elastic incoherent neutron scattering Increased flexibility associated with the inactivation of the oxygen evolving complex, Photosynth. Res., 111 (2012) 113–124.

[13] T. Matsuo, J. Peters, Sub-Nanosecond Dynamics of Pathologically Relevant Bio-Macromolecules Observed by Incoherent Neutron Scattering, Life, 12 (2022) 1259.

[14] P.J. Steinbach, R.J. Loncharich, B.R. Brooks, The effects of environment and hydration on protein dynamics: A simulation study of myoglobin, Chem. Phys., 158 (1991) 383–394.

[15] R. Guzzi, C. Arcangeli, A.R. Bizzarri, A molecular dynamics simulation study of the solvent isotope effect on copper plastocyanin, Biophys. Chem., 82 (1999) 9–22.

[16] P. Cioni, G.B. Strambini, Effect of Heavy Water on Protein Flexibility, Biophys. J., 82 (2002) 3246–3253.

[17] P. Sasisanker, A. Oleinikova, H. Weingärtner, R. Ravindra, R. Winter, Solvation properties and stability of ribonuclease A in normal and deuterated water studied by dielectric relaxation and differential scanning/pressure perturbation calorimetry, Phys. Chem. Chem. Phys., 6 (2004) 1899–1905.

[18] J.B. Linse, J.S. Hub, Three- And four-site models for heavy water: SPC/E-HW, TIP3P-HW, and TIP4P/2005-HW, J. Chem. Phys., 154 (2021).

[19] M.J. Abraham, T. Murtola, R. Schulz, S. Páll, J.C. Smith, B. Hess, E. Lindahl, GROMACS: High performance molecular simulations through multi-level parallelism from laptops to supercomputers, SoftwareX, 1–2 (2015) 19–25.

[20] Y. Duan, C. Wu, S. Chowdhury, M.C. Lee, G. Xiong, W. Zhang, R. Yang, P. Cieplak, R. Luo, T. Lee, J. Caldwell, J. Wang, P. Kollman, J.C. Chem, A point-charge force field for molecular mechanics simulations of proteins based on condensed-phase quantum mechanical calculations, J Comput Chem, 24 (2003) 1999–2012.

[21] W.L. Jorgensen, J. Chandrasekhar, J.D. Madura, R.W. Impey, M.L. Klein, Comparison of simple potential functions for simulating liquid water, J. Chem. Phys., 79 (1983) 926–935.

[22] C.P. Lawrence, J.L. Skinner, Flexible TIP4P model for molecular dynamics simulation of liquid water, Chem Phys Lett, 372 (2003) 842–847.

[23] R. Agarwal, M.D. Smith, J.C. Smith, Capturing Deuteration Effects in a Molecular Mechanics Force Field: Deuterated THF and the THF-Water Miscibility Gap, J. Chem. Theory Comput., 16 (2020) 2529–2540.

[24] P.G. Hill, R.D. MacMillan, V. Lee, Tables of thermodynamic properties of heavy water in SI units, Canada, 1981.

[25] M. Bée, Quasielastic Neutron Scattering, Adam Hilger, Bristol and Philadelphia, 1988.

[26] S. Dellerue, A.J. Petrescu, J.C. Smith, M.C. Bellissent-Funel, Radially softening diffusive motions in a globular protein, Biophys. J., 81 (2001) 1666–1676.

[27] G. Williams, D.C. Watts, Non-symmetrical dielectric relaxation behaviour arising from a simple empirical decay function, Trans. Faraday Soc., 66 (1970) 80–85.

[28] K. Singwi, A. Sjölander, Diffusive Motions in Water and Cold Neutron Scattering, Phys. Rev., 119 (1960) 863–871.

[29] F. Volino, A.J. Dianoux, Neutron incoherent scattering law for diffusion in a potential of spherical symmetry: general formalism and application to diffusion inside a sphere, Mol. Phys., 41 (1980) 271–279.

[30] R.G.J. Miller, Simultaneous Statistical Inference, Springer Science & Business Media, 2012.

[31] F. Merzel, J.C. Smith, Is the first hydration shell of lysozyme of higher density than bulk water?, Proc Natl Acad Sci U S A, 99 (2002) 5378–5383.

[32] A. Arbe, J. Colmenero, M. Monkenbusch, D. Richter, Dynamics of glass-forming polymers: “Homogeneous” versus “heterogeneous” scenario, Phys. Rev. Lett., 81 (1998) 590–593.

[33] S. Herrig, M. Thol, A.H. Harvey, E.W. Lemmon, A Reference Equation of State for Heavy Water, J. Phys. Chem. Ref. Data, 47 (2018) 43102.

[34] P.A. Baghurst, L.W. Nichol, W.H. Sawyer, The effect of D2O on the association of β-lactoglobulin A, J. Biol. Chem., 247 (1972) 3199–3204.

[35] T.J. Itoh, H. Sato, The effects of deuterium oxide (2H2O) on the polymerization of tubulin in vitro, BBA - Gen. Subj., 800 (1984) 21–27.

[36] H. Omori, M. Kuroda, H. Naora, H. Takeda, Y. Nio, H. Otani, K. Tamura, Deuterium oxide (heavy water) accelerates actin assembly in vitro and changes microfilament distribution in cultured cells, Eur. J. Cell Biol., 74 (1997) 273–280.

[37] G. Chakrabarti, S. Kim, M.L. Gupta, J.S. Barton, R.H. Himes, Stabilization of tubulin by deuterium oxide, Biochemistry, 38 (1999) 3067–3072.

[38] K. Pounot, H. Chaaban, V. Foderà, G. Schirò, M. Weik, T. Seydel, Tracking Internal and Global Diffusive Dynamics During Protein Aggregation by High-Resolution Neutron Spectroscopy, J. Phys. Chem. Lett., 11 (2020) 6299–6304.

[39] T. Matsuo, A. De Francesco, J. Peters, Molecular Dynamics of Lysozyme Amyloid Polymorphs Studied by Incoherent Neutron Scattering, Front. Mol. Biosci., 8 (2022) 812096.

[40] T. Matsuo, T. Tominaga, F. Kono, K. Shibata, S. Fujiwara, Modulation of the picosecond dynamics of troponin by the cardiomyopathy-causing mutation K247R of troponin T observed by quasielastic neutron scattering, Biochim Biophys Acta Proteins Proteom, 1865 (2017) 1781–1789.

